# Coarse-Grained Dynamic Simulation of Cytoskeletal Microtubule Twist

**DOI:** 10.1101/2024.10.16.618782

**Authors:** Renjie Zhu, Yuwei Zhang, Fei Xia

## Abstract

Microtubule, a key component of the cytoskeleton, plays a crucial role in cell mitosis. Modeling and dynamic simulation of microtubules at the micrometer scale remain a significant challenge. In this study, we developed the helix-based ultra-coarse-grained (HB-UCG) model for the dynamic simulation of microtubules, based on the helical characteristics obtained from electron microscopy density data. We constructed microtubules up to 35 μm in length and investigated the relationship between persistence length and contour length. By comparing our results with experimental data, we validated the potential of the HB-UCG model in simulating microtubule functions. We also simulated the twist process of microtubules driven by motor proteins, successfully demonstrating the model’s effectiveness in simulating biological processes. This study can provide the foundation for theoretical simulations of microtubule functions during mitosis.

## 1 Introduction

Microtubules (MTs) are widely distributed in eukaryotic cells, playing critical biological roles^1^. As one of the essential components of the cytoskeleton^2^, microtubules participate in intracellular material transport and maintain the shape and functional integrity of cells. During mitosis, microtubules are generated by the centrosomes to form spindle fibers, promoting nuclear division to generate two new daughter cells. Given the importance of microtubule function, researchers have employed a variety of experimental and theoretical approaches to explore the biomechanical properties of microtubules at the nanoscale, observing their involvement in crucial cellular processes, such as microtubule polymerization-depolymerization dynamics^3^, motor protein movement along microtubules^4^, and the chiral features generated during mitosis^5^. Ivec et al. observed that microtubule bundles forming the spindle underwent directional twisting with a certain helicity during cell mitosis^6^. Recently, Mitra et al. reported in their study of motor protein-driven microtubule twisting that single microtubule, under the influence of kinesin-14, wrap around each other in an antiparallel configuration, forming multiple turns^4^. Their findings contribute to a deeper understanding of the biological functions of microtubule bundles during mitosis. Therefore, elucidating the helicity and chiral properties of individual microtubules is important for comprehending the role of spindle fibers in mitotic processes.

Early experiments conducted numerous measurements on the biomechanical properties of single microtubules, focusing primarily on parameters such as the Young’s modulus *E*, shear modulus *G*^7, 8^, bending stiffness *κ*^9-11^, and persistence length *l*_p_^12, 13^. For instance, using techniques such as thermal fluctuations^14^, optical tweezers^15^, and atomic force microscopy^16^, the Young’s modulus *E* has been reported to range between 0.31 and 7.0 GPa^17^. Pampaloni and coworkers employed single-particle tracking methods to measure persistence lengths of microtubules ranging from 2.6 to 47.5 μm^13^. Based on the experimental data, they fitted a nonlinear equation describing the relationship between persistence length and the contour length, discovering that as the measured contour length increases, the persistence length *l*_p_ approaches an asymptotic value 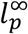. Experimental results indicate that the mechanical properties of microtubules, such as persistence length *l*_p_, display a dependence on the length of the contour length. However, there are certain limitations to these experimental measurements, such as limitations of the maximum growth length of microtubules, making it difficult to obtain data for microtubules longer than 20 μm.

Theoretical simulations provide an important complementary approach for studying the functional properties of microtubules. However, there are at least two major challenges in simulating microtubules. The first challenge lies in the fact that the true scale of microtubules is in the micron range, with approximately 42 million atoms per micron, which makes all-atom (AA) simulations computationally unattainable. For example, Tong et al. reported an AA simulation of a 64 nm microtubule for 500 ns^18^, which still falls far short of the real biological scale. The second challenge is that microtubules are hollow cylindrical structures formed by helical arrangements of tubulins, displaying anisotropic mechanical properties that cannot be adequately described using classical elastic models such as the worm-like chain model^19, 20^. To accurately simulate the mechanical properties of microtubules, theoretical models should not only capture the correct structural features of the microtubule, but also scale up to the micron-level macroscopic dimensions.

To enable dynamic simulations of microtubules at the micron scale, coarse-grained (CG) models of microtubules have been developed and reported in the literature. For instance, Deriu et al. constructed an elastic network model for a 350 nm microtubule^21^, while Dima et al. developed a SOP coarse-grained model for microtubules^22^. Recently, our group developed an ultra-coarse-grained (UCG) model for microtubules ranging from 1 to 12 μm in length, based on low-resolution cryo-electron microscopy data of microtubules^23^. Using the UCG models, we simulated and analyzed the relationship between the persistence length *l*_p_ and the contour length of microtubules, attaining results that were consistent with experimental findings^24^. Nevertheless, the length of microtubules in cells often exceeds 12 μm, requiring the development of models that can simulate longer microtubules to fully capture their functions and properties in cellular environments.

In this study, we developed a helix-based ultra-coarse-grained (HB-UCG) model for microtubules, based on the helical characteristics of the microtubule structure. Using the HB-UCG model, we simulated the persistence length and the dynamic process of twist formation in microtubules ranging from 12 to 35 μm in length. This research provides an important theoretical approach for elucidating the functions of microtubules during mitosis in cells through simulation.

## 2 Results

We first introduce the potential energy function and parameterization process of the HB-UCG model, which is a coarse-grained model for microtubules within the length range of 2 to 35 μm. To validate the accuracy of the HB-UCG model, we performed long-timescale coarse-grained molecular dynamics (CGMD) simulations to calculate the persistence length *l*_*p*_ of microtubules ranging from 12 to 35 μm, and we successfully predicted the asymptotic persistence length 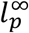. After confirming the validity of the model, we further applied it to replicate the dynamic process of microtubule twisting, as observed in experimental studies.

### 2.1 Development of MT HB-UCG Models

Inspired by the characteristic helical arrangement of tubulin during microtubule polymerization, we developed the UCG models based on the helical features of microtubules. The modeling process is illustrated in **Figure 1**. The modeling data were derived from low-resolution cryo-electron microscopy (cryo-EM) measurements of a 32 nm microtubule (EMD-4043)^25^, as shown in **Figure 1a**. The original cryo-EM data provided three-dimensional coordinate information of the density volume. To reduce noise from smaller density data, we set a threshold of 240, and further extracted the region beyond 11.6 nm from the center using a higher threshold of 300, obtaining the helical distribution of both internal and external density layers (**Figure 1b**). Next, we performed a weighted averaging of the spatial coordinates based on the density (**Figure 1c**), and fitted the helix equation (**Figure 1d**) **eq.(1):**

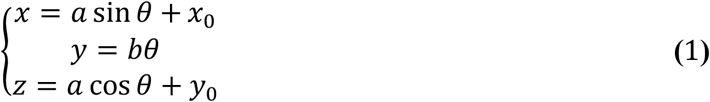

where *a* represents the radius of the circular cross-section of the helix, and *x*_*0*_ & *y*_*0*_ are the coordinates of the center of the circle. The parameter *b* characterizes the pitch of the helical structure for both inner and outer helices. Based on the helix equations, a series of UCG beads are generated based on the characteristic that the microtubule consists of 13 protofilaments^2^. Computational details can be found in section **4.1**. Compared to the previously established UCG representation using the convolutional and K-means coarse-graining (CK-CG) method^24^, the advantage of the HB-UCG model lies in its utilization of the regularity in the helical density arrangement, allowing the representation of microtubules of arbitrary length. This avoids the need for additional coarse-graining optimization of the density data. In this study, the HB-UCG model we developed can simulate microtubules up to 35 μm in length, whereas the longest microtubule that could be modeled using the CK-CG method is 12 μm.

**Figure 1:**
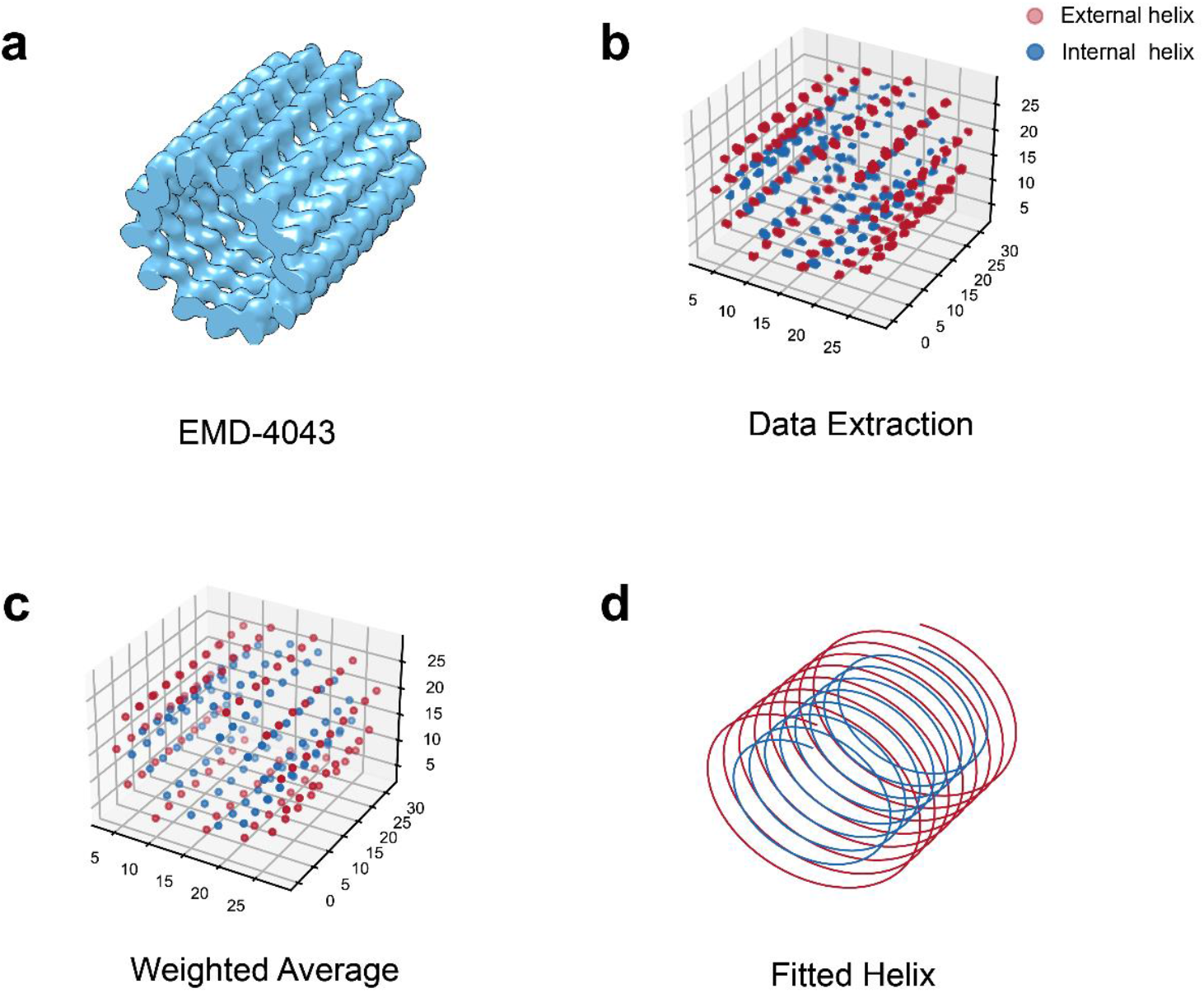
HB-UCG model construction process. (**a**) The low-resolution cryo-electron microscopy (cryo-EM) data of the microtubule EMD-4043; (**b**) The initial data for fitting is extracted; (**c**) Weighted averaging is conducted to obtain data points for fitting the internal/external helix equations; (**d**) The helix obtained from the fitting can be used to generate UCG beads.

As shown in **Figure 2a**, the HB-UCG beads are categorized into two types based on their position within the microtubule structure: internal and external UCG beads. By comparing the HB-UCG representation with the corresponding microtubule structure, it can be concluded that a single tubulin monomer is primarily represented by one external and one internal UCG bead. The average molar mass of a tubulin monomer is approximately 55000 g/mol^26^. Based on the ratio of the densities between internal and external distribution in the electron microscopy data, we deduce that the mass of the internal and external UCG beads are 25715 g/mol and 29249 g/mol, respectively.

**Figure 2:**
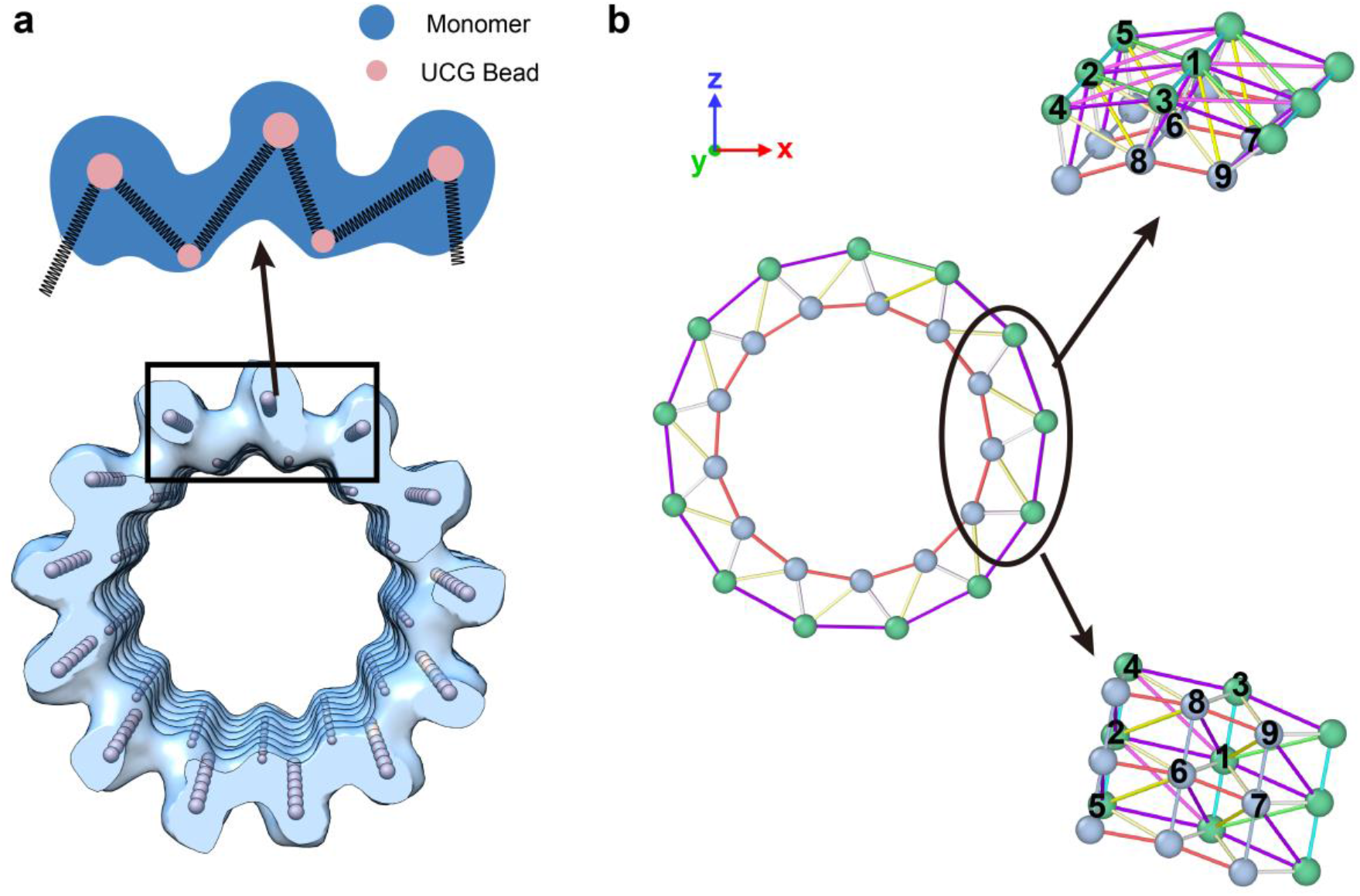
Schematic representation of monomers and UCG beads and the bonding method between UCG beads. **(a)** Superimposed view of UCG beads and microtubule cryo-electron microscopy density map observed from a cross-sectional perspective; **(b) Left**: A cross-sectional view showing the method of bond addition between UCG beads. **Right**: A portion of the HB-CG model from two different angles demonstrating the bond addition method. Different colored bonds represent different types of bonding methods, and different colored beads represent different types of UCG beads. The numerical labels correspond to the detailed bond descriptions listed in **Table 1**.

The interaction environment of internal and external UCG beads determines that there can be up to 10 different bonding connections between any two adjacent UCG beads, as depicted in **Figure 2b**. The bonding interactions between any two UCG density beads are described using an anisotropic elastic network model (MVP-ANM)^27^. In the MVP-ANM model, the bonding interaction between any two UCG beads *i* and *j*, with a distance *d*_*ij*_, is represented by a spring with an elastic constant *k*_*ij*_, and the equilibrium distance between the beads is denoted as 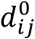. The total potential energy of the entire HB-UCG model system can be described by **eq.(2)**:

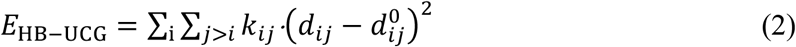

**Table 1.** Bonds formed between UCG beads and the parameters of potential energy function in eq.(2).

| **Bonds** | **$k_{ij}$ (N/m)** | **$d_{ij}^0$ (nm)** |
| --- | --- | --- |
| 1-2 | 0.03044 | 5.866 |
| 1-3 | 0.09676 | 3.989 |
| 1-4 | 0.009668 | 7.264 |
| 1-5 | 0.01313 | 6.919 |
| 6-7 | 0.06423 | 4.283 |
| 6-8 | 0.07479 | 3.989 |
| 1-7 | 0.05613 | 4.750 |
| 1-6 | 0.09770 | 3.707 |
| 1-9 | 0.02303 | 6.068 |
| 1-8 | 0.034392 | 5.514 |

The specific expression for the elastic force constant *k*_*ij*_ is given in **eq.(3)**

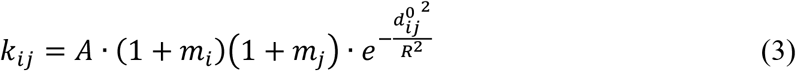

where *k*_*ij*_ depends on the masses of beads *i* and *j*, denoted as *m*_*i*_ and *m*_*j*_. The empirical parameter *R* reflects the magnitude of heterogeneous interactions, and in this study, it is set to 4 nm. The parameter *A* is a scaling factor, determined by fitting the experimental values of the Young’s modulus.

As shown in **Figure 3a**, we constructed a 200 nm long microtubule HB-UCG model and performed 200 ns of tensile dynamics simulations. By fitting the experimental data of the Young’s modulus at 300K, with a reported value of 100 MPa^28^, we determined the empirical parameter *A* to be 1.1×10^41^ kg^-1^·s^-2^. Further details of the tensile dynamic simulations can be found in the section **4.2**.

**Figure 3:**
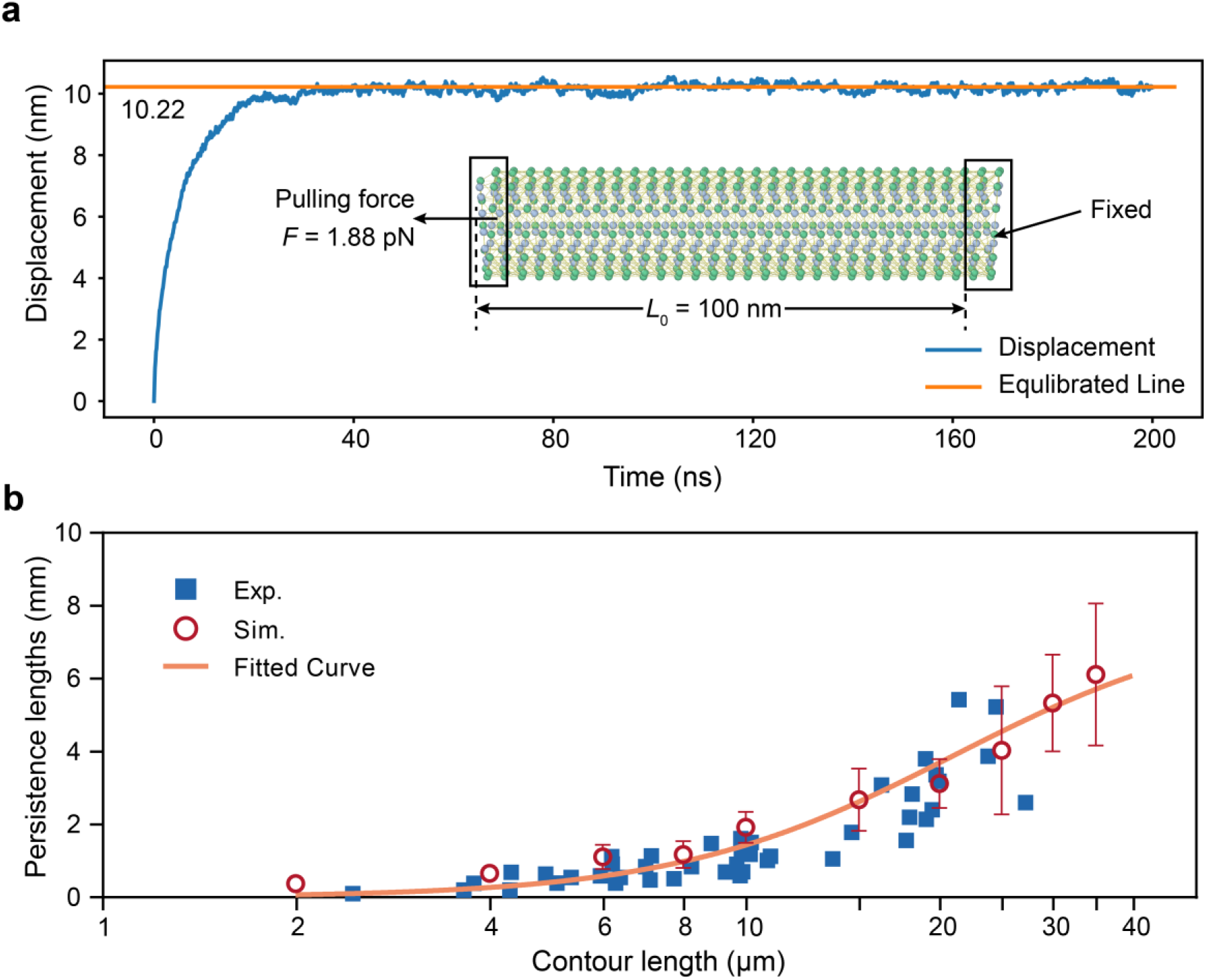
Results and methods for young’s modulus measurement and the relationship between microtubule persistence length and contour length. **(a)** Schematic of the method for determining the Young’s modulus of the microtubule. One 100 nm end is fixed, and a constant force is applied to a 4 nm section at the top of the free end, causing displacement until equilibrium is reached. The average displacement of the free end of microtubule gradually stabilizes during the simulation (**blue curve**), with the final equilibrium displacement being 10.22 nm (**orange line**); **(b)** The blue solid squares represent previous experimental results^13^, while the red hollow circles represent the simulation results, with five parallel simulations conducted for each contour length. The orange curve shows the fitting results using **eq.(2)**.

### 2.2 CGMD Simulations of Persistence Lengths

Persistence length is one of the key mechanical properties of biological materials, reflecting the material’s resistance to bending. Pampaloni et al. used thermal fluctuation techniques to measure the persistence lengths of microtubules ranging from 2 to 35 μm in length^13^. Based on the measured persistence length *l*_*p*_ of microtubules with a contour length *L*, they proposed that as the microtubule contour length increases, the persistence length approaches a constant value 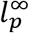. The relationship between *l*_*p*_ and 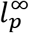 is described by **eq.(4)**:

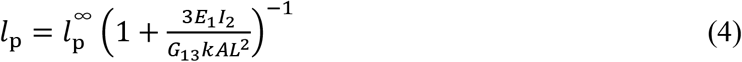

In their experiments, Pampaloni et al. reported that most microtubules measured were shorter than 20 μm. Due to limitations in experimental conditions affecting microtubule growth, data points for microtubules in the 20-35 μm range are scarce. In our previous coarse-grained molecular dynamics (CGMD) studies^24^, we successfully simulated the experimental values of for microtubules up to 12 μm in length. However, due to limitations in theoretical models and computational resources, it was hard to simulate microtubules longer than 12 μm.

In this study, our newly developed HB-UCG model allows for the simulation of longer microtubules. We extended the simulated length of microtubules up to 12-35 μm and performed millisecond-level CGMD simulations for microtubules of varying lengths. We selected 10 different lengths, *L* = 2, 4, 6, 8, 10, 15, 20, 25, 30 and 35 μm, for modeling and calculations. For each length, we performed five parallel CGMD simulations, each lasting 1.2 ms. We computed the average persistence length and the standard deviations for microtubules of each fixed length, and the results are shown in **Table 2**.

**Table 2.** CGMD simulation results for the persistence lengths of MTs whose contour lengths *L*_*0*_ range from 2 to 35 μm.

| $L_0$ ($\mu\text{m}$) | Persistence Lengths ($\mu\text{m}$) |
| --- | --- |
| | Average Values ± SD |
| 2.0 | 381.0 ± 127.5 |
| 4.0 | 659.4 ± 169.4 |
| 6.0 | 1115 ± 334.7 |
| 8.0 | 1175 ± 364.7 |
| 10.0 | 1921 ± 422.4 |
| 15.0 | 2680 ± 857.4 |
| 20.0 | 3123 ± 672.1 |
| 25.0 | 4035 ± 1759 |
| 30.0 | 5335 ± 1332 |
| 35.0 | 6117 ± 1948 |

**Figure 3b** presents a comparison between the statistical results from our CGMD simulations and the experimental values. From this, it is clear that the experimental measurements for microtubules shorter than 25 μm are relatively comprehensive, and our simulation results, including the error ranges, show a good agreement with the experimental data within this length range. For microtubules with lengths *L* = 30 and 35 μm, the average *l*_*p*_ calculated from our simulations were 5.3±1.3 mm and 6.1±1.9 mm, respectively. Due to the increased fluctuations in longer microtubules, the standard deviations reached 1.3 mm and 1.9 mm. We fitted the simulated data from **Table 2** using **eq.(3)** and the value of 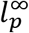 is 7.8 mm, which is consistent with the range reported by Pampaloni et al., which is 6.3±0.8 mm^13^. These CGMD simulations of single microtubules demonstrate that our developed HB-UCG model can accurately capture the mechanical properties of single microtubules. This provides a solid foundation for further studies on simulating the torque and dynamic behaviors of microtubules.

### 2.3 CGMD Simulation of Microtubule Twist

Recently, Mitra et al. reported in cell-based experiments that the motion of motor proteins along microtubules leads to torsional deformation and the formation of turns^4^. This significant experiment can reveal that the motion of motors can induce condition changes in microtubules. Motivated by these findings, we aim to utilize our developed HB-UCG model to simulate the dynamic process of twist formation at the molecular level, thereby elucidating the underlying mechanical properties and mechanisms involved in this process. During the formation of twist, interactions between different sections of the microtubule structure play a crucial role. To prevent the formation of unphysical spatial interactions between UCG beads in different parts of the microtubule, we introduced a repulsive interaction term^29^ into **eq.(2)** of the HB-UCG model, as described in **eq.(5)**:

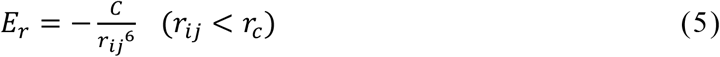

Where *r*_*ij*_ represents the distance between UCG beads *i* and *j*, and *r*_*c*_ is the cutoff radius. The empirical values of the parameters *C* and *r*_c_ are 2.08×10^51^ kg·s^-2^·m^-4^ and 3.5 nm, respectively.

In our CGMD simulation (details provided in the section **4.4**), we mimicked the experimental conditions, where one end of the microtubule was locked and the other end was continuously rotated by external forces (**Figure 4a**). As shown in **Figure 4b**, as the simulation progressed, the first turn formed at the simulation time of 383 μs, then followed by the formation of the second turn at about 708 μs, and the third turn was nearly complete by 803 μs, as shown in **Figure 4c**. The CGMD simulation resulted in the formation of three turns on a 10 μm spatial scale and near-millisecond timescale. As illustrated in **Figure 4b**, we selected the top 1000 UCG beads from the simulation trajectory and used the least squares method to fit the equation of the plane where these beads exist and calculate the normal vector of the plane. The angle between this normal vector and the z-axis was computed and recorded. **Figure 4d** shows the periodic changes of this angle over time during the simulation process, confirming the progression of the twisting behavior.

**Figure 4.**
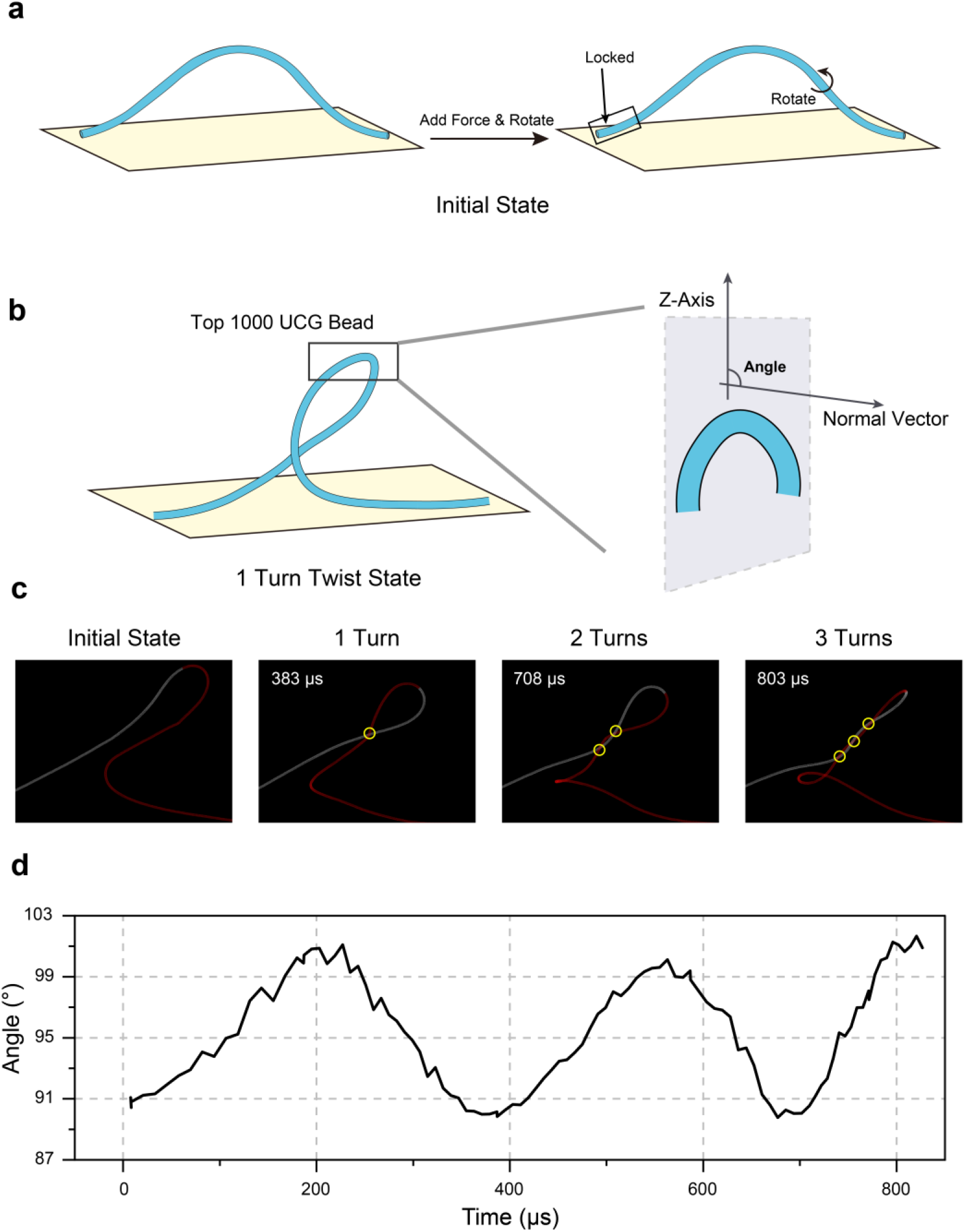
Simulation method and results of microtubule twist. **(a) Left**: Initial state of the microtubule before the application of torque (displayed in an arched shape); **Right**: Simulation strategy simulating motor protein movement along the microtubule. In reference to the experimental method by Mitra et al.^4^, one end is fixed while the other end is rotated to the left to apply torque; **(b)** Illustration showing how the angle value on the left axis of the line graph in **(d)** is calculated; **(c)** From left to right, four images showing the progression of twisting in a 10 μm microtubule as the simulation proceeds. At 803 μs, three turns have formed. The red color indicates the movable end, and the white color indicates the fixed end; **(d)** Graph showing how the angle changes over time as the twisting process progresses.

## 3 Discussion

In this study, we developed the HB-UCG model for microtubules based on the helix features observed in cryo-electron microscopy density data, with the capability to simulate microtubules up to 35 μm in length. We parameterized the HB-UCG model and simulated the persistence lengths of microtubules ranging from 2 to 35 μm. From the simulation trajectories, we calculated the asymptotic persistence length 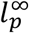 to be 7.8 mm, which is consistent with the range reported by Pampaloni et al. (6.3 ± 0.8 mm)^13^, thus validating the accuracy of the HB-UCG model.

Additionally, we employed the HB-UCG model to simulate the dynamic process of twist formation in a single microtubule under external forces, shedding light on the relationship between the changes in the mechanical properties of microtubules and their functions in the presence of actin. Furthermore, our model can serve as a foundation for multiscale modeling, where dynamic simulations of multiscale systems can be used to explore how conformational changes during microtubule dynamics influence their functionality^30^.

## 4 Methods

### 4.1 Coarse-graining Density based on Helix Equations

First, we set a density threshold and extracted the density data from cryo-electron microscopy, resulting in a distribution exhibiting helical characteristics. The distribution of the density data reflects the morphology of the internal and external density distribution of the tubulin within the microtubule. The determination of the density threshold was carried out using an empirical method, where a series of thresholds were tested to identify the optimal value that best captured these features.

After data extraction, we obtained a cluster of density points in three-dimensional coordinates. We selected the density values as weights and performed a weighted average of the coordinates for all points within a cluster to obtain the centroid of that cluster. The series of centroids obtained were then used for following fitting calculations. The weighted average coordinates were then fitted using the helix equation **eq.(1)**.

The key parameters of the helix are its radius and pitch. We first projected all points onto the *x-z* plane and applied the least squares method to fit the parameters *a, x*_0_ and *y*_0_ in the equations for the internal and external helices. Once these three parameters were determined, we projected all points onto the *x-y* and *y-z* planes and applied the least squares method twice to fit and determine the average value of *b*. The final fitted equations for the internal and external helices are as follows (units: nm): Inner helix equation is **eq.(6)**:

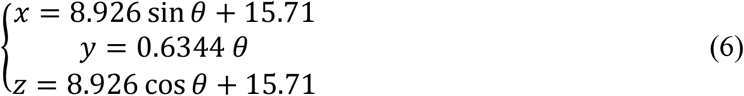

Outer helix equation is **eq.(7)**:

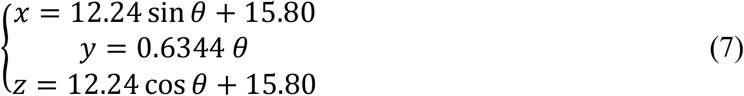

### 4.2 Model Parameterization and Simulation Details

For determining the value of the parameter *R* in **eq.(3)**, we used the LAMMPS software package^31^ to simulate a 1 μm microtubule. The entire simulation system was set up under the NVT ensemble, and a Langevin thermostat was used to maintain a constant temperature of 300 K. Under non-periodic boundary conditions, the relaxation time for the dynamic simulation was set to 10 ps, the integration time step for the simulation was set to 3 ps, and the total simulation duration was 0.1 ms. Ultimately, once the system stabilized, the value of *R* was determined to be 4 nm. For the determination of the parameter *A* in **eq.(3)**, we conducted tensile CGMD dynamics simulations of the microtubule and fitted the experimental data for the Young’s modulus *E*_exp_ which is 100 MPa^28^. From this fitting process, the value of the parameter *A* was derived to be 1.1×10^41^ kg^-1^·s^-2^. Specifically, we selected a 200 nm microtubule model, with one end fixed, and applied a constant force of 1.88 nN to the UCG beads at the other end, stretching the microtubule axially. The simulation ensemble and temperature conditions were the same as previously described, with an integration time step of 10 fs and a total simulation duration of 200 ns. The Young’s modulus *E*_cal_ was calculated using **eq.(8)**^32^. In this equation, *F* represents the applied constant force, *S* is the cross-sectional area of the microtubule, Δ*L* is the displacement of the movable end of the microtubule (as shown in **Figure 3a**), and *L*_0_ is the initial length of the movable end of the microtubule.

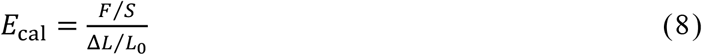

### 4.3 CGMD Simulation of Persistence Lengths

The CGMD simulation conditions for persistence length simulation are the same as previously described, with a total simulation time of 1.2 ms. The calculation of the persistence length *l*_*p*_ is based on **eq.(9)**:

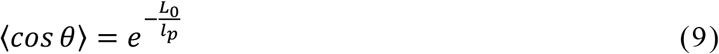

where *L*_*0*_ represents the contour length of the microtubule, and ⟨cos*θ*⟩ means the average value of the cosine of θ (0 ≤ θ ≤ π/2). The angle *θ* denotes the intersection angle of two tangent lines at the positions of the head and tail of microtubules. To calculate this, the coordinates of UCG beads within 40 nm from both ends of the microtubule are used. 3D linear regression is then applied to fit a straight line for each end, and the angle θ between these two lines is measured.

### 4.4 CGMD Simulation of Formation of MT Twists

In the simulation process, we used the LAMMPS software package to simulate a 10 μm long microtubule. The simulation system was conducted under the NVT ensemble, with a Langevin thermostat used to maintain a constant temperature of 300 K. The relaxation time for the dynamic simulation was set to 10 ps, and the integration time step was set to 1.5 ps under non-periodic boundary conditions. Initially, we selected the middle section of the microtubule for stretching, which deformed the microtubule into an arched shape, as shown in **Figure 4a**. Following the experimental observation method^4^, one end of the microtubule was fixed, while the other end was continuously rotated, simulating motor protein activity. This can be implemented in LAMMPS by using the fix move rotate command. A 0.5 μm section at the movable end was rotated around the central axis with a period of 100 μs. The simulation continued until multiple-turns structure appeared, as depicted in **Figure 4c**.

## Acknowledgements

This work was supported by the National Natural Science Foundation of China (No. 22073029, 22403051) and the National Undergraduate Training Programs for Innovation and Entrepreneurship (No. 202310269048G). We also acknowledge the support of the NYU-ECNU Center for Computational Chemistry at NYU Shanghai as well as the ECNU Public Platform for Innovation (001) for providing computer time.

